# Personalized logical models to investigate cancer response to BRAF treatments in melanomas and colorectal cancers

**DOI:** 10.1101/2020.05.27.119016

**Authors:** Jonas Béal, Lorenzo Pantolini, Vincent Noël, Emmanuel Barillot, Laurence Calzone

## Abstract

The study of response to cancer treatments has benefited greatly from the contribution of different omics data but their interpretation is sometimes difficult. Some mathematical models based on prior biological knowledge of signalling pathways, facilitate this interpretation but often require fitting of their parameters using perturbation data. We propose a more qualitative mechanistic approach, based on logical formalism and on the sole mapping and interpretation of omics data, and able to recover differences in sensitivity to gene inhibition without model training. This approach is showcased by the study of BRAF inhibition in patients with melanomas and colorectal cancers who experience significant differences in sensitivity despite similar omics profiles.

We first gather information from literature and build a logical model summarizing the regulatory network of the mitogen-activated protein kinase (MAPK) pathway surrounding BRAF, with factors involved in the BRAF inhibition resistance mechanisms. The relevance of this model is verified by automatically assessing that it qualitatively reproduces response or resistance behaviours identified in the literature. Data from over 100 melanoma and colorectal cancer cell lines are then used to validate the model’s ability to explain differences in sensitivity. This generic model is transformed into personalized cell line-specific logical models by integrating the omics information of the cell lines as constraints of the model. The use of mutations alone allows personalized models to correlate significantly with experimental sensitivities to BRAF inhibition, both from drug and CRISPR targeting, and even better with the joint use of mutations and RNA, supporting multi-omics mechanistic models. A comparison of these untrained models with learning approaches highlights similarities in interpretation and complementarity depending on the size of the datasets.

This parsimonious pipeline, which can easily be extended to other biological questions, makes it possible to explore the mechanistic causes of the response to treatment, on an individualized basis.

**Author summary:** We constructed a logical model to study, from a dynamical perspective, the differences between melanomas and colorectal cancers that share the same BRAF mutations but exhibit different sensitivities to anti-BRAF treatments. The model was built from the literature and completed from existing pathway databases. The model encompasses the key proteins of the MAPK pathway and was made specific to each cancer cell line (100 melanoma and colorectal cell lines from public database) using available omics data, including mutations and RNAseq data. It can simulate the effect of drugs and show high correlation with experimental results. Moreover, the structure of the network confirms both the importance of the reactivation of the MAPK pathway through CRAF and the involvement of PI3K/AKT pathway in the mechanisms of resistance to BRAF inhibition.

The study shows that, because of the low number of samples, the mechanistic approach that we propose provides different insights than powerful standard machine learning methodologies would, showing the complementarity between the two approaches. An important aspect to mention is that the mechanistic approach presented here does not rely on training datasets but directly interprets and maps data on the model to simulate drug responses.

## Introduction

In the age of high-throughput sequencing technologies, cancer is considered to be a genetic disease for which driver genes are constantly being discovered [1]. The study of mutational and molecular patterns in cancer patients aims to improve the understanding of oncogenesis. However, many of these gene alterations seem to be specific to certain cancer types [2] or exhibit different behaviours depending on the molecular context, particularly in terms of response to treatment. This prompted a shift from univariate biomarker-based approaches to more holistic methodologies leveraging the various omics data available.

To study these observed differences in drug response in various cancers, some approaches based on mathematical modelling were developed to explore the complexity of differential drug sensitivities. A number of machine learning-based methods for predicting sensitivities have been proposed [3], either without particular constraints or with varying degrees of prior knowledge; but they do not provide a mechanistic understanding of the response. Some other approaches focused on the description of the processes that might influence the response by integrating knowledge of the signalling pathways and their mechanisms and translated it into a mathematical model [4–6]. The first step of this approach implies the construction of a network recapitulating knowledge of the interactions between selected biological entities (driver genes but also key genes of signalling pathways), extracted from the literature or from public pathway databases, or directly inferred from data [7]. This static representation of the mechanisms is then translated into a dynamical mathematical model with the goal to not only understand the observed differences [5] but also to predict means to revert unexpected behaviours.

One way to address issues related to patient response to treatments is to fit these mechanistic models to the available data, and to train them on high-throughput cell-line specific perturbation data [4, 5, 8]. These mechanistic models are then easier to interpret with regard to the main drivers of drug response. They also enable the *in silico* simulations of new designs such as combinations of drugs not present in the initial training data [6].

However, these mechanistic models contain many parameters that need to be fitted or extracted from the literature. Some parsimonious mathematical formalisms have been developed to make up for the absence of either rich perturbation data to train the models or fully quantified kinetic or molecular data to derive the parameters directly from literature. One of these approaches is the logical modelling, which uses discrete variables governed by logical rules. Its explicit syntax facilitates the interpretation of mechanisms and drug response [9, 10] and despite its simplicity, semi-quantitative analyses have already been performed on complex systems [11] for both cancer applications [9, 12] and drug response studies [13, 14], and have proved their efficacy [15, 16].

The nature of this formalism has shown its relevance in cases where the model is not automatically trained on data but simply constructed from literature or pathway databases and where biological experiments focus on a particular cell line [17]. We propose here a pipeline based on logical modelling and able to go from the formulation of a biological question to the validation of a mathematical model on pre-clinical data, in this case a set of cell lines (Fig 1), and the subsequent interpretation of potential resistance mechanisms. The application of the mechanistic model to different cell lines is therefore done without any training of parameters but only on the basis of the automatic integration and interpretation of their omic features.

**Fig 1.**
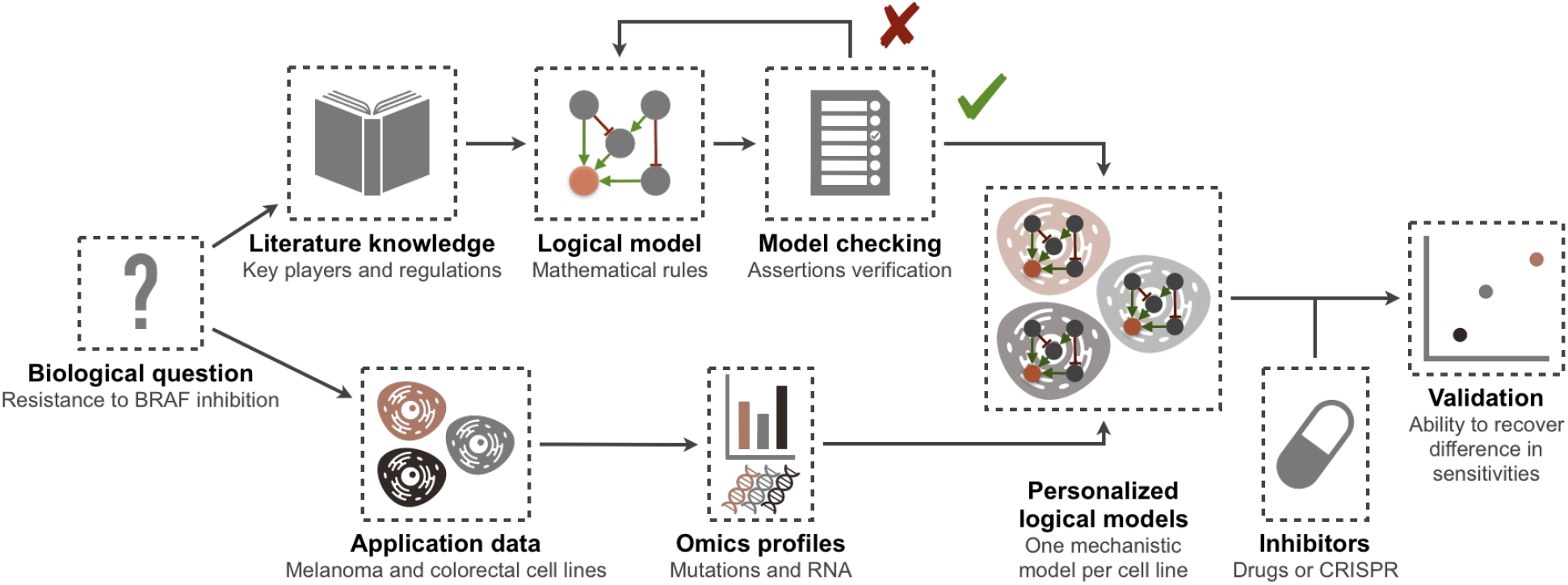
BRAF modelling flowchart: from a biological question to validated personalized logical models.

The construction of a mathematical model must be based first and foremost on a precise and specific biological problem, at the origin of the design of the model. Here, we choose to explore the different responses to treatments in diverse cancers that bear the same mutation. A well-studied example of these variations is the BRAF mutation and especially its V600E substitution. BRAF is mutated in 40 to 70% of melanoma tumours and in 10% of colorectal tumours, each time composed almost entirely of V600E mutations [18]. In spite of these similarities, BRAF inhibition treatments have experienced opposite results with improved survival in patients with melanoma [19] and significant resistance in colorectal cancers [20], suggesting drastic mechanistic differences. Some subsequent studies have proposed context-based molecular explanations, often highlighting surrounding genes or signalling pathways, such as a feedback activation of EGFR [21] or other mechanisms [22, 23]. These various findings support the need for an integrative mechanistic model able to formalize and explain more systematically the differences in drug sensitivities depending on the molecular context. The purpose of the study we propose here is not to provide a comprehensive molecular description of the response but to verify that the existence and functionality of the suggested feedback loops around the signalling pathway in which BRAF is involved [21] may be a first hint towards these differences. For a more thorough study of these cancers, we refer to other works [4, 24, 25].

A logical model summarizing the main molecular interactions at work in colorectal cancers and melanomas is thus built from the literature and completed with databases. As previously mentioned, the objective is to understand whether it is possible to model and explain differences in responses to BRAF inhibition in melanoma and colorectal cancer patients using the same regulatory network. The fact that the two cancers share the same network but differ from the alterations and expression of their genes constitute our prior hypothesis. We then use model checking tools to verify the consistency of this model with a series of qualitative assertions retrieved from literature. Finally, we use available public omics data from these cancer cell lines to transform the generic model into personalized cell-line models. The relevance of the latter is validated by their ability to recover the differences in BRAF inhibition sensitivities, from both drug and CRISPR screenings.

## Materials and methods

### Logical model principles and simulations

A concise overview of the main properties of logic modelling is provided and additional details may be found in dedicated reviews [26, 27]. A logical model can be represented by a regulatory graph where nodes are biological entities and edges are influences of one entity onto the others. Each node is considered as a discrete variable (0, 1 or more if required) corresponding to the activity level of the associated biological entity (0 is inactive or absent, 1 is active or present) (Fig 2A). Each entity (proteins, genes, etc.) can thus represent distinct biological states (e.g., expressed gene, phosphorylated protein, etc.) depending on the meaning that is given to each of the nodes. These entities are connected by edges representing positive (resp. negative) influences, i.e., activation (resp. inhibition) of the target node by the sourced node. Combinatorial outcome of influences on one node is defined by the logical rules assigned to the node and expressed with logical operators AND (&), OR (|) and NOT (!), as in Fig 2A. The dynamics of this mathematical model can be expressed using the state transition graph (STG) where the nodes of this graph represent the states of the model. In the STG, the edges show the possible transitions between the different model states according to the logical rules (Fig 2B). As these rules often allow for different transitions, either all of them can be performed at each time step (synchronous update) or performed sequentially by choosing how the priorities are defined (asynchronous updates) [26, 27]. For the rest of the article, the term “node” will refer to those of the regulatory network.

**Fig 2.**
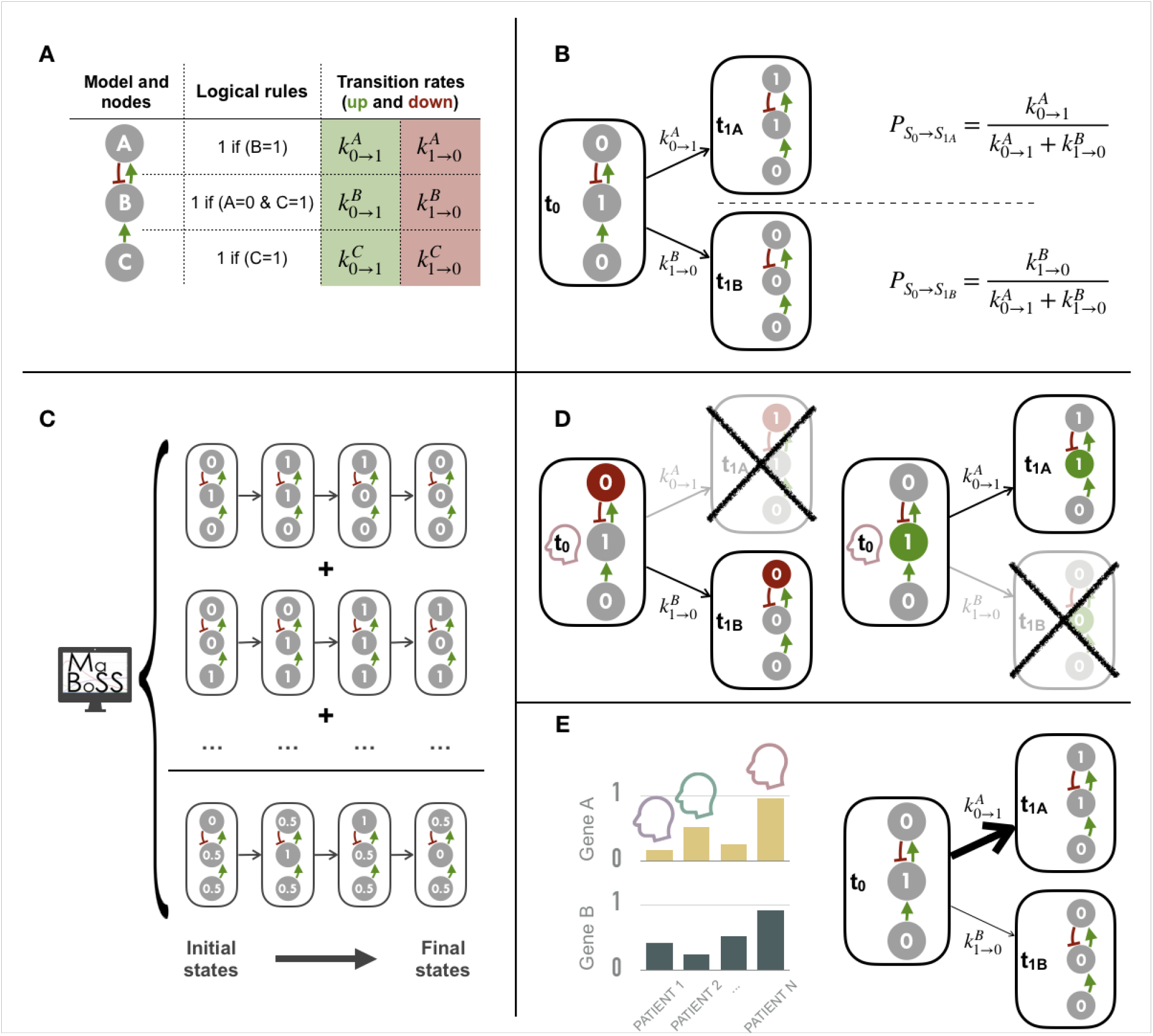
Logical modelling: principles, simulation and personalization. (A) A simple example of a logical model with three nodes: the regulatory graph, the corresponding logical rules and the transition rates as used in MaBoSS [28]. (B) State transition graph of the logical model with the two possible transitions resulting from the given initial conditions and the probabilities of choosing stochastically one of them. (C) Schematic representation of a logic model simulation with MaBoSS: average trajectory obtained from the mean of many individual stochastic trajectories. (D) Personalization of a logical model with discrete data: a node forced to stay at 0, on the left (resp. 1 on the right) prevents (resp. favours) some transitions from occurring. (E) Personalization of a logical model with continuous data: since gene A is highly expressed in the red patient, the probability of activation of the corresponding node is increased (resp. probability of inhibition is decreased for gene B)

Note that a variable of the model is considered to be active or inactive depending on the biological meaning that is assigned to the node. The active state will differ according to the nature of the entity: genes and proteins may be active if they are expressed. If some proteins are known to be kinases, their targets will be considered to be active (i.e., their value goes from 0 to 1) in their phosphorylated form, in most cases. The choice will be made for each individual node of the network. That way, BRAF, CRAF, MEK, ERK, AKT, GAB1, MDM2 and p70 are considered to be active in their phosphorylated form. When SOX10 is phosphorylated by ERK, it can no longer mediate the synthesis of FOXD3. In this case, the active form is the unphosphorylated one. It is translated in the model by a direct inhibitory influence of ERK on SOX10 target.

In the present article, all simulations are performed according to asynchronous updating with MaBoSS software [11, 28]. This algorithm, using continuous time Markov chain simulations on the Boolean network, provides a stochastic way to choose a specific transition among several possible ones. Each node is associated with transition rates, either for activation of the node *k*_0→1_ (or *k_up_*) or inactivation *k*_1→0_ (or *k_down_*) and the stochastic choice between the possible transitions is made based on these transition rates (Fig 2B). For our simulations, unless otherwise specified (cf. section about personalization of models), all transition states were initially assigned to 1. The exploration of all the state space of the model is then done by simulating a very large number of individual stochastic trajectories in order to aggregate them into a mean stochastic trajectory (Fig 2C). To ensure a proper exploration of the state space, the number of computed stochastic trajectories should increase with the model complexity. In the present work, all simulations are performed with 5000 trajectories after verifying that this number was sufficient to ensure very low variability in the final results. 100 different simulations of the generic model, with 5000 trajectories each, result in an average *Proliferation* score of 0.182 with a standard deviation of 0.005 across the 100 simulations.

The scores obtained after each simulation correspond to the final asymptotic states, i.e., the average stochastic state reached by the model after a defined period of time. *t_end_* = 50 was chosen because at this time, it was ensured that the solutions had reached their asymptotic state by comparing with values reached at later times (average *Proliferation* score of 0.182 also at *t_end_* = 100 with 100 simulations of 5000 trajectories).

### Automatized model-checking within unit-testing framework

The construction of a model is a daunting task, as each improvement is susceptible to change the dynamical properties of the model. To tackle this problem, we need a simple way to test these properties and detect if the model is still able to reproduce them. Software development knows similar challenges, where improvements can break existing functionalities. Software verification thus became an important part of software development, which assess whether a software meets a list of requirements. After each important modification, tests are run to verify that the software still produces the expected behaviour. A similar framework can be applied for model construction to check the validity of the model for each iteration of the building process. First, the modeller must describe what is the expected behaviour of the model for several conditions, based either on scientific literature or biological experiments. Some similar works using model checking to build and verify models, were recently published [29, 30].

In order to standardize this process, we developed a tool, called MaBoSS_test, to easily verify if a logical model was coherent with specific biological assertions. This tool is based on MaBoSS simulation software, which produces simulations describing the evolution of the probability of states with time. Inspired from python’s unittest library, we developed an extension for MaBoSS simulations which tests the validity of the dynamical behaviour of the model via assert methods. Each assertion is used to verify if the model satisfies a given type of biological statement.

The majority of the tests consist in altering the model, by changing the initial condition or introducing a set of mutations, then observing how these alterations affect the probability of reaching a specific state with respect to the original simulation. An example of a biological assertion may be the reactivation of the MAPK (mitogen-activated protein kinase) pathway through EGFR signal after BRAF inhibition in colorectal cancer [21]. To test if our model is consistent with this statement, we call the function:

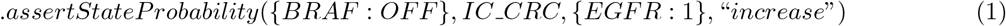

The arguments of this method are the following: the set of mutations to perform (BRAF:OFF), the predefined initial conditions of the simulation (IC_CRC), the state we wish to observe (EGFR:1), and a string to specify the behaviour (increase). The function can be read as: “Assert that: after BRAF inhibition, using the initial condition for the colorectal cancer, ***IC_CRC***, the probability of EGFR activation increases”. If the “unit testing output” is selected, in the case that the assertion is not correct, the test will fail raising an exception. Otherwise, if the “detailed output” is selected, the result will be “True” or “False” and the probabilities to have EGFR active before and after BRAF inhibition will be displayed. This tool, its documentation and an example in the form of a Jupyter notebook are available on GitHub (https://github.com/sysbio-curie/MaBoSS_test).

### Cell lines omics profiles

The omics profiles of colorectal and melanoma cell lines are downloaded from Cell Model Passports portal [31]. 64 colorectal cancer (CRC) cell lines and 65 cutaneaous melanoma (CM) cell lines are listed in the database, with at least mutation or RNA-seq data (59 CM and 53 CRC with both mutations and RNA-seq data).

### Personalization of logical models with cell lines omics data

The PROFILE (PeRsonalization OF logIcaL ModEls) methodology transforms a generic logical model into as many personalized models as there are cell lines by using and integrating their omics profiles [32]. The general idea is to rely on the interpretation of the omics data and translate it into constraints of the mathematical model.

The method to integrate omics data are separated into two strategies: for discrete data (i.e., mutations, copy number alterations) and for continuous data (i.e., transcriptomics, (phospho)proteomics). The discrete strategy consists in setting the value of a node to 0 or 1 for the whole duration of the simulation. In the present work, this is done based on mutation data and functional effect inference. The mutations identified in the cell lines are interpreted using OncoKB database [33], an evidence-based repository with mutation annotations. Mutations referenced as loss-of-function (resp. gain-of function) are forced to 0 (resp. 1), thus constraining the possible transitions in the model as in Fig 2D, left (resp. right). Uninterpreted mutations, which are by far the majority, are not included in the models. The distribution of mutations in the four most frequently mutated genes is shown in Fig 3A.

**Fig 3.**
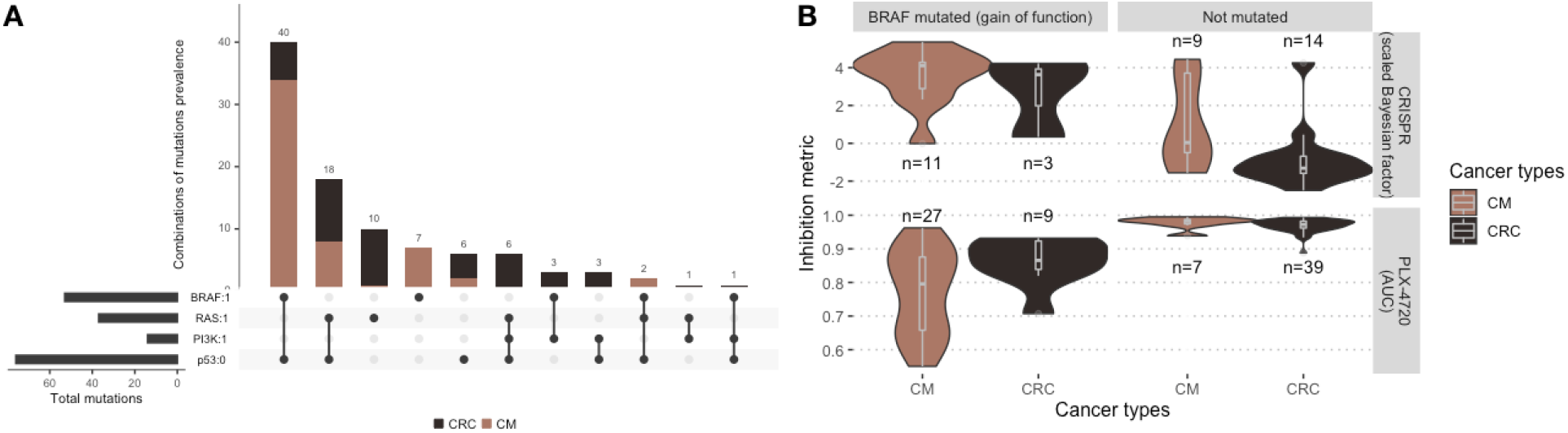
Cell lines data: mutations and sensitivities to BRAF inhibition. (A) Distribution of the assigned mutations in the four most frequently mutated genes in the colorectal/melanoma cohort of cell lines [34]. (B) Differential sensitivities to BRAF inhibition by the drug PLX-4720 (lower panel) or by CRISPR inhibition (upper panel), depending on BRAF mutational status and cancer type. Numbers of cell lines in eache category are indicated. Note that high sensitivities correspond to low AUC and high scaled Bayesian factors.

The second strategy, preferably used with continuous data, is to modify the initial conditions and transition rates based on a continuous proxy for node activity. A cell line with a clear over-expression of a gene/protein, compared to the whole cohort of interest, will have the transition rate related to the activation (resp. inhibition) of the corresponding node favoured and made more (resp. less) likely (Fig 2E). The initial probability that the node will be activated (i.e., the probability to start at 1 among the 5000 stochastic trajectories) will also be modified accordingly, which is particularly important for input nodes that will not be regulated by the model. This method requires different conditions. First, a relevant node activity proxy has to be available in the data: it can be a level from transcriptomics, proteomics or even phospho-proteomics. Then, unlike mutations, the interpretation is not made absolutely but only in comparison to the other members of the cohort. In the present work, despite the proteic nature of most of the model nodes, only RNA data is available and is therefore used, on the assumption that it can be a consistent, although not ideal proxy, as it has sometimes been proposed in other studies [35]. Note that the distribution of RNA levels is normalized between 0 and 1 on a gene-specific basis before being included in the model.

### Drug and CRISPR/Cas9 screening

In order to validate the relevance of personalized models to explain differential sensitivities to drugs, some experimental screening datasets are used. Drug screening data are downloaded from the Genomics of Drug Sensitivity in Cancer (GDSC) dataset [36] which includes two BRAF inhibitors: PLX-4720 and Dabrafenib. The cell lines are treated with increasing concentration of drugs and the viability of the cell line relative to untreated control is measured. The dose-response relative viability curve is fitted and then used to compute the half maximal inhibitory concentration (IC50) and the area under the dose-response curve (AUC) [37]. Since the IC50 values are often extrapolated outside the concentration range actually tested, we will focus on the AUC metric for all validation with drug screening data. AUC is a value between 0 and 1: values close to 1 mean that the relative viability has not been decreased, and lower values correspond to increased sensitivity to inhibitions. The results obtained with the two drugs are very strongly correlated (Pearson correlation of 0.91) and the analyses presented here will therefore focus on only one of them, PLX-4720.

Results of CRISPR/Cas9 screening are downloaded from Cell Model Passports [31]. Two different datasets from Sanger Institute [38] and Broad Institute [39] are available. We use scaled Bayesian factors to assess the effect of CRISPR targeting of genes. These scores are computed based on the fold change distribution of sgRNA [40]. The highest values indicate that the targeted gene is essential to the cell fitness. The agreement between the two databases is good [41] but we choose to focus on the Broad database, which is more balanced in terms of the relative proportions of melanomas and colorectal cancers.

Fig 3B illustrates both the relative quantities of cell lines for which drug or CRISPR screening data are available (depending on their BRAF status) as well as differences in sensitivity to BRAF inhibition. The greater sensitivity of BRAF-mutated melanomas compared to BRAF-mutated colorectal cancers is well observed for PLX-4720. However, the overlap in the distributions requires a deeper look into the data and a search for more precise explanations of the differences in sensitivity, including within each type of cancer. The finding appears to be similar for CRISPR despite a sample size that is too small; the higher average sensitivity of melanomas even extends to non-mutated BRAF.

### Validation of personalized models using CRISPR/Cas9 and drug screening

The validation of personalized logical models using these screening data is done with the following rationale. First, the models are personalized using omics data from the cell lines. Then, two separate simulations are performed for each personalized model: one without the inhibition, the other by creating and activating a BRAF inhibitor to mimic the drug or CRISPR inhibition. The ratio of the *Proliferation* phenotype obtained with inhibition and without inhibition is the proxy used to be compared with the different screening metrics each of which is also standardized (AUC calculated on relative viability for drugs and Bayes factor computed from fold-changes and then scaled).

### Random forests

Random forests are used as an example of a machine learning approach to compare with mechanistic models [42] and are implemented with *randomForestSRC* R package. Random forests can be seen as an aggregation of decision trees, each trained on a different training set formed by uniform sampling with replacement of the original cohort. Prediction performances are computed using out-of-the bag estimates for each individual (i.e., average estimate from trees that did not contain the individual in their bootstrap training sample) and summarized as percentage of variance explained by the random forest. It is also possible to compute the variable importance that assesses the contribution of variables to the overall performance. The solution adopted in this paper to measure it, and called VIMP in the package, consists in introducing random permutations between individuals for the values of a variable and quantifying the variation in performance resulting from this addition of noise. In the case of key variables for prediction, this perturbation will decrease the performance and will result in a high variable importance [43].

## Results

### A generic logical model for melanoma and colorectal cancers

The construction of the logical model aims at summarizing the current molecular understanding of BRAF gene and its molecular partners in both colorectal cancers and melanomas. The focus of this model is put on two important signalling pathways involved in the mechanisms of resistance to BRAF inhibition which are the ERK1/2 MAPK and PI3K/AKT pathways [44, 45].

The MAPK pathway encompasses three families of protein kinases: RAF, MEK, ERK. If RAF is separated into two isoforms, CRAF and BRAF, the other two families MEK and ERK are represented by a single node. When BRAF is inhibited, ERK can still be activated through CRAF, and BRAF binds to and phosphorylates MEK1 and MEK2 more efficiently than CRAF [46], especially in his V600E/K mutated form. When PI3K/AKT pathway is activated, through the presence of the HGF (Hepatocyte Growth Factors), EGF (Epidermal Growth Factors) and FGF (Fibroblast Growth Factors) ligands, it leads to a proliferative phenotype. The activation of this pathway results in the activation of PDPK1 and mTOR, both able to phosphorylate p70 (RPS6KB1) which then promotes cell proliferation and growth [47]. There has been some evidence of negative regulations of these two pathways carried out by ERK itself [48]: phosphorylated ERK is able to prevent the SOS-GRB2 complex formation through the activation of SPRY [49], inhibit the EGF-dependent GAB1/PI3K association [50] and down-regulate EGFR signal through phosphorylation [48]. The model also accounts for a negative regulation of proliferation through a pathway involving p53 activation in response to DNA damage (represented by ATM); p53 hinders proliferation through the activation of both PTEN, a PI3K inhibitor, and p21 (CDKN1A) responsible for cell cycle arrest.

The generic network presented in Fig 4 recapitulates the known interactions between the biological entities of the network. The network was first built from the literature, and then was verified and completed with potential missing connections using SIGNOR database [51]. More details about the model can be found in the GINsim annotation file of the model [52], available in Supporting Information.

**Fig 4.**
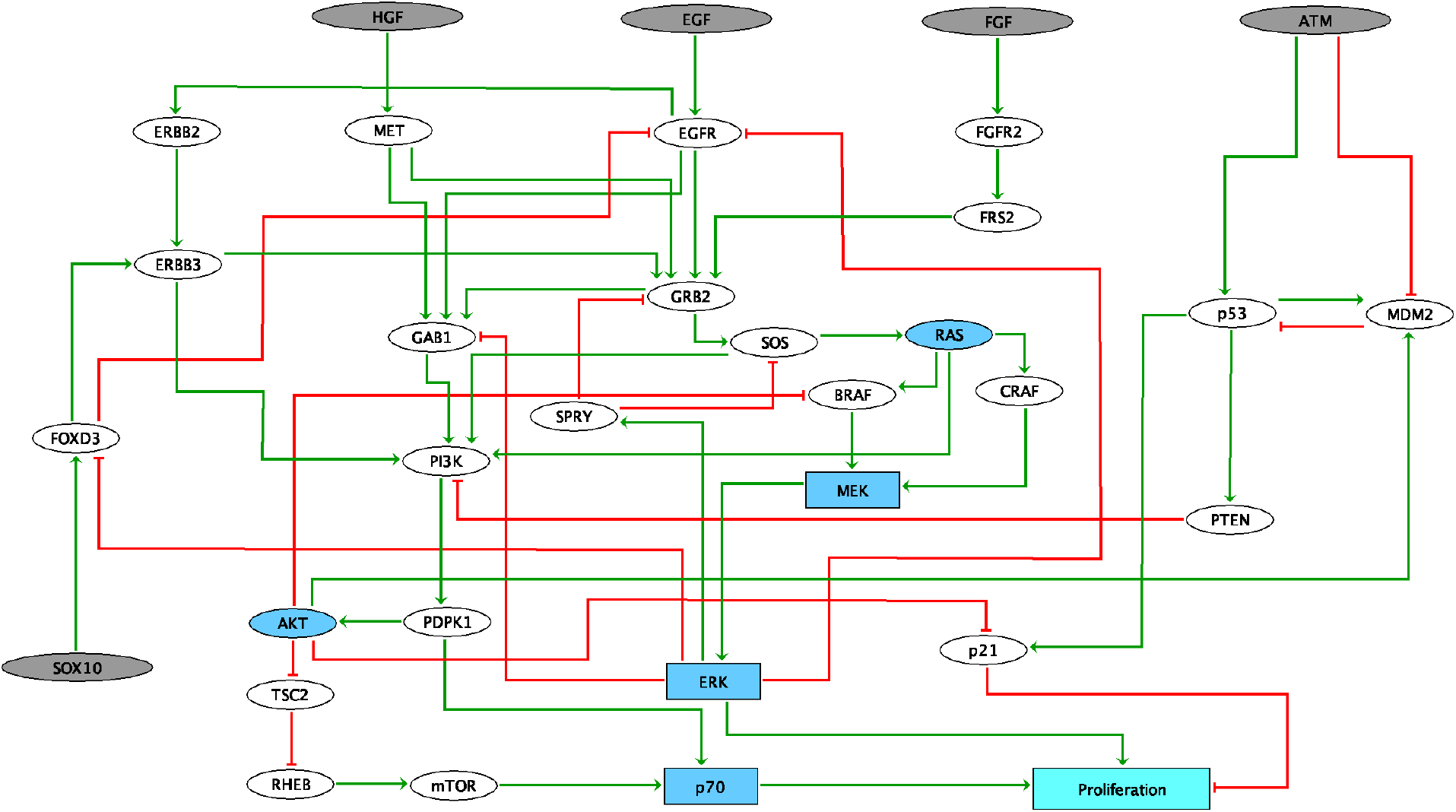
Logical model of signalling pathways around BRAF in colorectal and melanoma cancers. Grey nodes represent input nodes, which may correspond to the environmental conditions. Blue nodes accounts for families. Light blue node represents the output of the model. Square nodes represent multi-valued variable (MEK, ERK, p70 and Proliferation). Note that *Proliferation* is used as the phenotypic read-out of the model.

We hypothesize that a single network is able to discriminate between melanoma and CRC cells. These differences may come from different sources. One of them is linked to the negative feedback loop from ERK to EGFR. As mentioned previously, this feedback leads to one important difference in response to treatment between melanoma and CRC: *BRAF*^(*V*600*E*)^ inhibition causes a rapid feedback activation of EGFR, which supports continued proliferation. This feedback is observed only in colorectal since melanoma cells express low levels of EGFR and are therefore not subject to this reactivation [21]. Moreover, phosphorylation of SOX10 by ERK inhibits its transcription activity towards multiple target genes by interfering with the sumoylation of SOX10 at K55, which is essential for its transcriptional activity [53]. The absence of ERK releases the activity of SOX10, which is necessary and sufficient for FOXD3 induction. FOXD3 is then able to directly activate the expression of ERBB3 at the transcriptional level, enhancing the responsiveness of melanoma cells to NRG1 (the ligand for ERBB3), and thus leading to the reactivation of both MAPK and PI3K/AKT pathways [53]. Furthermore, it has been shown that in colorectal cells, FOXD3 inhibits EGFR signal in *vitro* [54]. Interestingly, SOX10 is highly expressed in melanoma cell lines when compared to other cancer cells. In the model, we define SOX10 as an input because of the lack of information about the regulatory mechanisms controlling its activity. The different expression levels of SOX10 have been reported to play an important role in melanoma (high expression) and colorectal (low expression) cell lines.

The features and expected behaviours for both cancers were formulated as assertions of the model and verified at each step of the model construction (Table 1) through the automatized model-checking framework described in Methods. The tool enables to easily extend the model with new components, while ensuring that the constraints on which the model was built are maintained.

**Table 1.** List of assertions used to validate the logical model.

| Assertions | Refs |
| --- | --- |
| 1: *BRAF inhibition causes a feedback activation of EGFR in colorectal cancer and not in melanoma.* | [21] |
| 2: *MEK inhibition stops ERK signal but activates the PI3K/Akt pathway and increases the activity of ERBB3.* | [48, 55] |
| 3: *HGF signal leads to the reactivation of the MAPK and PI3K/AKT pathways, and resistance to BRAF inhibition.* | [56] |
| 4: *BRAF inhibition in melanoma activates the SOX10/FOXD3/ERBB3 axis, which mediates resistance through the activation of the PI3K/AKT pathway.* | [53] |
| 5: *Overexpression/mutation of CRAF results in constitutive activation of ERK and MEK also in the presence of a BRAF inhibitor.* | [57] [58] |
| 6: *Early resistance to BRAF inhibition may be observed in case of PTEN loss, or mutations in PI3K or AKT.* | [57] |
| 7: *Experiments in melanoma cell lines support combined treatment with BRAF/MEK + PI3K/AKT inhibitors to overcome resistance.* | [57] |
| 8: *BRAF inhibition (Vemurafenib) leads to the induction of PI3K/AKT pathway and inhibition of EGFR did not block this induction.* | [59] |
| 9: *Induction of PI3K/AKT pathway signaling has been associated with decreased sensitivity to MAPK inhibition.* | [59] |

The logical model formalizes the knowledge compiled from different sources. It highlights the role of SOX10, FOXD3, CRAF, PI3K, PTEN and of EGFR in resistance to anti-BRAF treatments. The purpose here is not to suggest new pathways that may be responsible for resistance but to formally confirm what has been suggested and support hand-waving explanations with a mathematical model. The model can be further used to simulate drug experiments and suggest conditions for which the treatment may be efficient or not. Adapting the generic model to each cancer type or cancer cell line will allow to search for the samples that do not respond to treatment, suggest the possible reasons for this resistance, but still focusing on the model components (Supplementary material, Fig S1).

### Differential sensitivities to BRAF targeting explained by personalized logical models

Once the logical model consistency has been validated, personalized models are generated for each cell line by integrating their interpreted genomic features directly as model constraints or parameters. Their sensitivity to BRAF inhibition is then compared to experimentally observed sensitivities (Fig 5). In all the following analyses, we focus on three different personalization strategies using: only mutations as discrete personalization (Fig 5A, upper row), only RNA as continuous personalization (Fig 5A, middle row) or mutations combined with RNA (Fig 5A, lower row). These choices reflect first of all the following *a priori*: mutations are much more drastic and permanent changes than RNA, whose expression levels are more subject to fluctuation and regulation. The objective is also to answer the following questions: What type of data is most likely to explain the differences in responses? Is it relevant to combine them? Fig 5 shows an example of the type of analyses possible with personalized models, zooming in more and more on the details from Fig 5A to Fig 5C.

**Fig 5.**
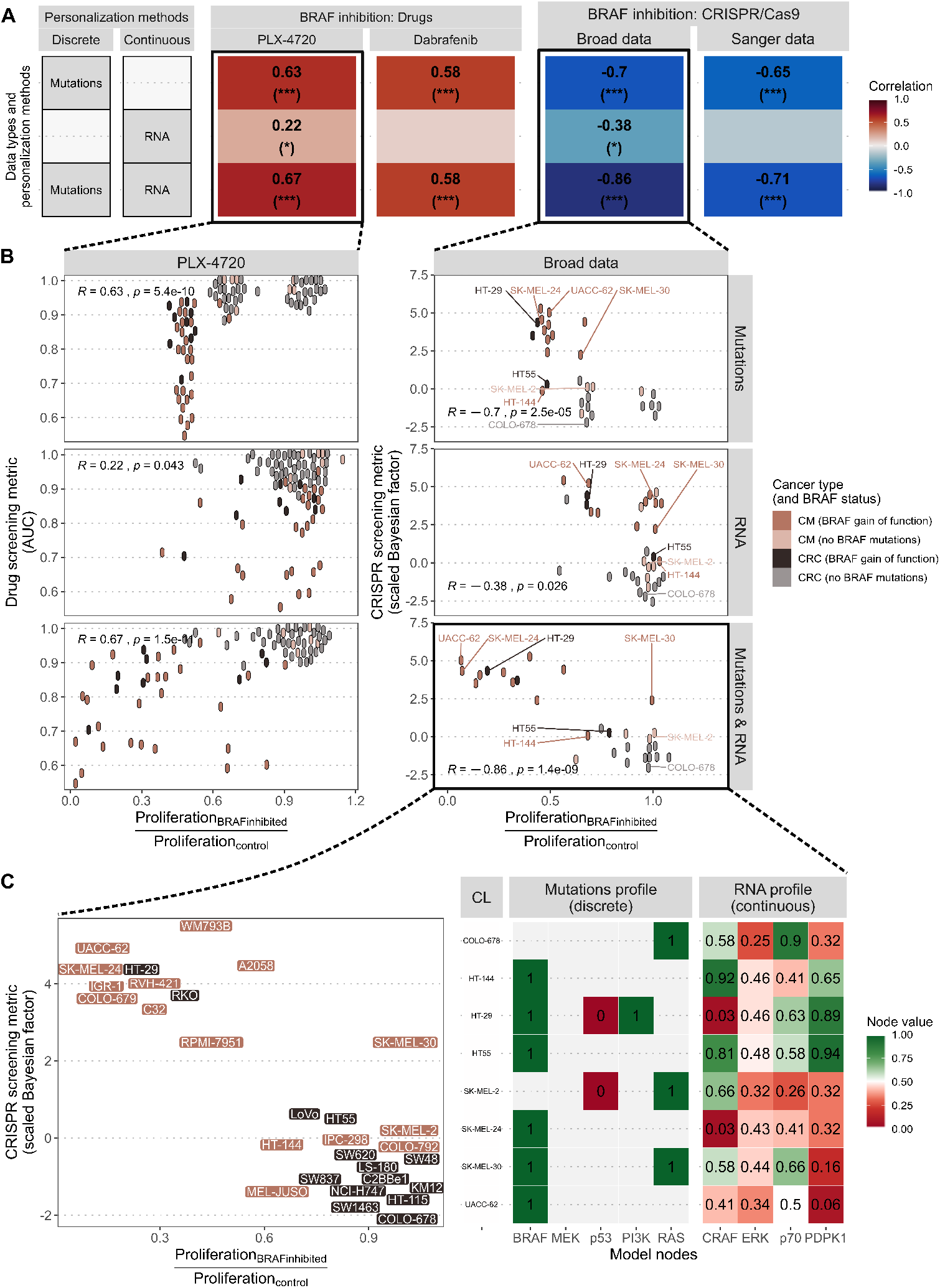
Validation of personalized models with cell lines data. (A) Pearson correlations between normalized *Proliferation* scores from personalized models and experimental sensitivities to BRAF inhibition by drug or CRISPR targeting; each row corresponds to a different personalization strategy; only the values for the significant correlations are displayed. (B) Scatter plots with non-overlapping points corresponding to correlations of panel A, with the three personalization strategies, focusing one one drug (PLX-4720) and one CRISPR dataset (Broad) only. (C) Enlargement of the scatter plot comparing model scores (personalized with mutations and RNA) and experimental sensitivity to CRISPR targeting of BRAF (left) with the corresponding table representing the omics profiles used for each cell line to explore the response mechanisms. This panel can be advantageously replaced by one of the interactive plots proposed in the provided code.

The first approach consists in using only mutations as discrete personalization (Fig 5A, upper row): the mutations identified in the dataset and that are present in the regulatory network are set to 1 for activating mutations and set to 0 for inactivating mutations. In this case, the *Proliferation* scores from personalized models significantly correlate with both BRAF drug inhibitors (PLX-4720 and Dabrafenib) and both CRISPR datasets (using Pearson correlations). Note that the opposite directions of the correlations for the drug and CRISPR datasets are due to the fact that cell lines sensitive to BRAF inhibition result in low AUCs, and high scaled Bayesian factors, respectively, and, if the models are relevant, to low standardized *Proliferation* scores. Looking more closely at the corresponding scatter plot for PLX-4720 (Fig 5B, upper left panel), it can be seen that this correlation results from the model’s ability to recover the highest sensitivity of the BRAF-mutated cell lines that form an undifferentiated cluster on the left side. These cell lines are indeed relatively more sensitive than non-mutated BRAF cell lines. However, the integration of mutations alone does not explain the significant differences within this subgroup (AUC between 0.55 and 0.9). A very similar behaviour can be observed when comparing model simulations with CRISPR data (Fig 5B, upper right panel).

Using only RNA data as continuous personalization (Fig 5A and B, middle rows) is both less informative and more difficult to interpret. For continuous data such as RNAseq data, we normalize the expression values, following the rules described in Methods section and in [32], and set both the initial conditions and the transition rates of the model variables to the corresponding values. Correlations with experimental BRAF inhibitions appear weaker and more uncertain.

The key point, however, is that the combination of mutations and RNA, as depicted in Fig 5A and B lower rows, seems to be more relevant. This is partially true in quantitative terms, looking at the correlation in Fig 5 but it is even easier to interpret in the corresponding scatter plots. Comparing first the Broad CRISPR scatter plots using mutations only (Fig 5B, upper right) and using both mutations and RNA (Fig 5B, lower right), we can observe that non-responsive cell lines (scaled Bayesian factor below 0), grouped in the lower right corner and correctly predicted using only mutations stayed in the same area: these strong mutational phenotypes have not been displaced by the addition of RNA data. Other cell lines previously considered to be of intermediate sensitivity by the model (e.g., COLO-678 or SK-MEL-2) were shifted to the right, consistent with the lack of sensitivity observed experimentally. Finally, BRAF-mutated cell lines, previously clustered in one single column on the left using only mutations (with normalized *Proliferation* scores around 0.5), have been moved in different directions. Many of the most sensitive cell lines (scaled Bayesian factor above 2) have been pushed to the left in accordance with the high sensitivities observed experimentally (e.g., HT-29 or SK-MEL-24). It is even observed that the model corrected the position of the two BRAF mutated cell lines, but whose sensitivity is experimentally low (melanoma cell line HT-144 and colorectal cell line HT-55). Only one cell line (SK-MEL-30) has seen its positioning evolve counter-intuitively as a result of the addition of RNA in the personalization strategy: relatively sensitive to the inhibition of BRAF, it has, however, seen its standardised *Proliferation* score approach 1. All in all, this contribution of RNA data results in significant correlations even when restricted to BRAF-mutated cell lines only (R=0.69, p.value=0.006).

A similar analysis can be made of the impact of adding RNA data to personalization when comparing with the experimental response to PLX-4720 (Fig 5B, upper and lower left panels). Most of the non-sensitive cell lines (upper right corner) have not seen the behaviour of the personalized models change with RNA addition. However, the numerous BRAF-mutated cell lines previously grouped around standardized *Proliferation* scores of 0.5, are now better differentiated and their sensitivity predicted by personalized models has generally been revised towards lower scores (i.e., higher sensitivity). Similar to the CRISPR data analysis, three sensitive cell lines have been shifted to the right and are misinterpreted by the model. As a result, the correlation restricted to BRAF-mutated cell lines is no longer significant (R=0.26, p.value=0.1).

### An investigative tool

These personalized models are not primarily intended to be predictive tools but rather used to reason and explore the possible mechanisms and specificities of each cell line. To continue on the previous examples, the two melanoma cell lines, HT-144 and SK-MEL-24, share the same mutational profiles but have very different sensitivities to BRAF targeting (Fig 5C). This inconsistency is partially corrected by the addition of the RNA data, which allows the model to take into account the difference in CRAF expression between the two cell lines. In fact, CRAF is a crucial node for the network since it is necessary for the reactivation of the MAPK pathway after BRAF inhibition. Therefore, the high sensitivity of SK-MEL-24 may be explained by its low CRAF expression level, which makes the reactivation of the MAPK pathway more difficult for this cell line. Conversely, in HT-144, the high level of CRAF expression allows the signal to flow properly through this pathway even after BRAF inhibition, thus making this cell line more resistant. The importance of CRAF expression is also evident in HT-29, a CRC BRAF mutated cell line with other important mutations (PI3K activation and p53 deletion). However, it remains sensitive to treatment, due to its very low level of CRAF expression.

Another interesting contribution of RNA appears in the melanoma cell line UACC-62, which is particularly sensitive to treatment. The model is able to correctly predict its response once RNA levels are integrated. In this case, the reason for sensitivity seems to be due to the low level of PDPK1, which makes it difficult to activate p70 and thus trigger the resistance linked to PI3K/AKT pathway activation. Similarly, the CRC resistant cell line, HT55, which carries only the BRAF mutation, expresses high levels of PDPK1, in addition to high levels of CRAF, supporting the idea that the presence of both MAPK and PI3K/AKT pathways may confer resistance to BRAF inhibition treatments. We can also mention a cluster of RAS mutated cell lines, usually NRAS mutated for melanomas (e.g., SK-MEL-2) and KRAS for colorectal cancers (e.g., COLO-678), which are classified by the model as resistant. Interestingly, in these cell lines, a low level of CRAF is not enough to block the signal of the MAPK pathway, which is stronger in the model because of the simulation of the RAS mutation (RAS is set to 1).

Only SK-MEL-30 appears to be incorrectly classified and is observed to be more sensitive than the other cell lines with a similar mutation profile. This could be due to the fact that our network is incomplete and not able to account for some alterations responsible for this cell line sensitivity. The exploration of the mutational profile for this cell line might be a hint of The problem may also come from the fact that this cell line contains a frameshift mutation of RPS6KB2 (p70 node) not referenced in OncoKB and therefore not included in the simulation.

The versatility of the logical formalism makes it possible to test other node inhibitions as in Fig 6, but remains limited by the scope of the model. Since the present model has been designed around BRAF, its regulators have been carefully selected and implemented, which is not necessarily the case for other nodes of the model. Therefore, these personalized models can be used to study how comprehensive the descriptions of the regulation of other nodes or parts of the model are. Thus, model simulations show that response trends to TP53 inhibition are consistently recovered by the model (Fig 6B) but the simple regulation of p53 in the model results in coarse-grained patterns, although slightly improved by addition of RNA data. Similar analyses regarding the targeting of PIK3CA (in CRISPR data) simulated, in the model, by the inhibition of PI3K node, can be performed (Fig 6C). Low correlations are an indication highlighting the insufficient regulation of the node.

**Fig 6.**
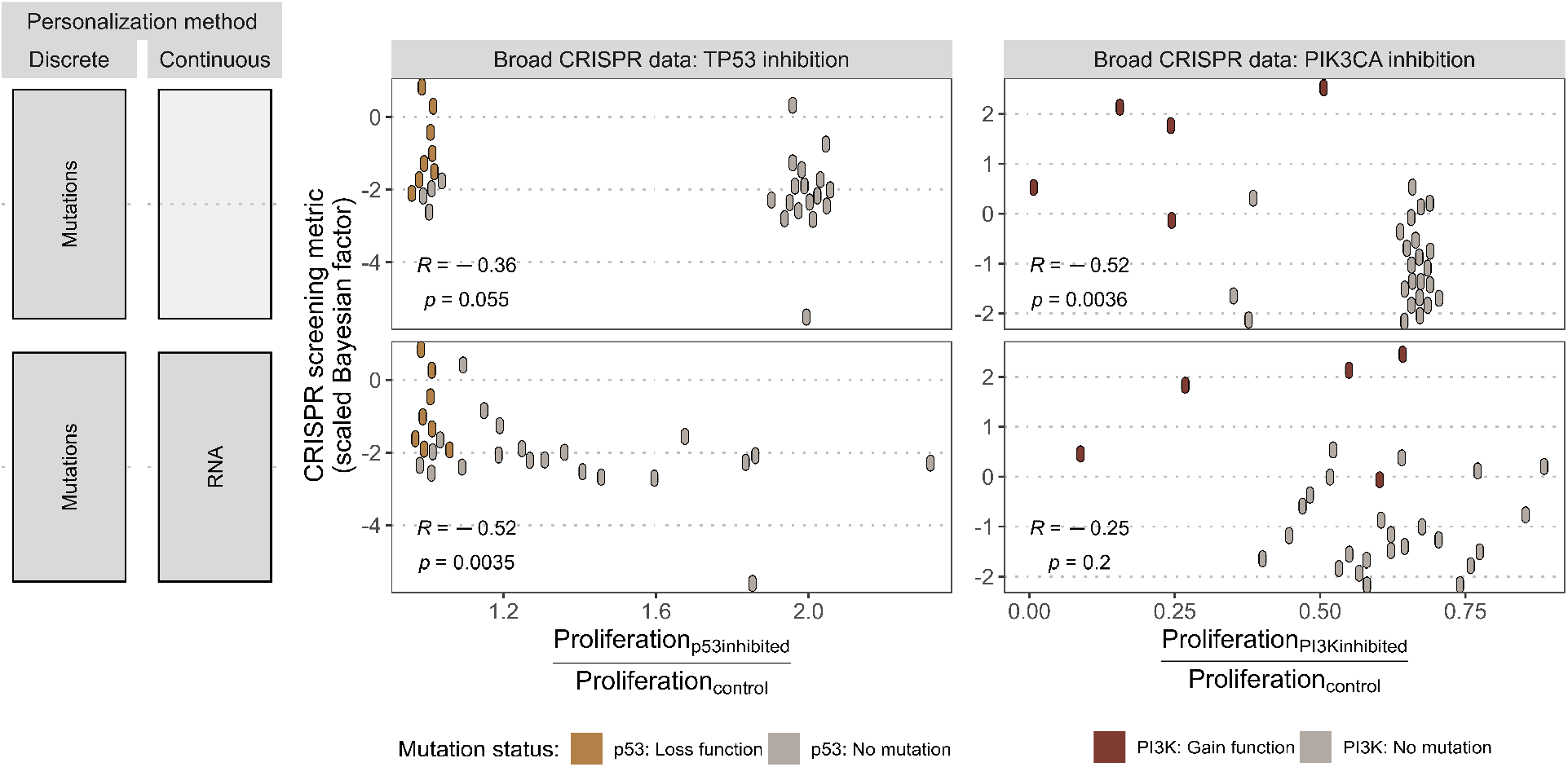
Application of personalized models to other CRISPR targets. (A) Personalization strategies using either mutations only (as discrete data) or combined with RNA (as continuous data) with their corresponding scatter plots in panels B and C. (B) Scatter plot comparing normalized *Proliferation* scores of p53 inhibition in the models with experiment sensitivity of cell lines to TP53 CRISPR inhibition, indicating p53 mutational status as interpreted in the model. Pearson correlations and the corresponding p-values are shown. (C) Similar analysis as in panel B with PI3K model node and PIK3CA CRISPR inhibition.

### Comparison of the mechanistic approach with machine learning methods

In order to provide comparison elements unbiased by prior knowledge or by the construction of the model, we performed some simple machine learning algorithms. Random forests have been fitted with inputs (mutations and/or RNA data) and outputs (sensitivities to drug or CRISPR BRAF inhibition) similar to those of logical models and the corresponding predictive accuracies are reported in Fig S2. The first insight concerns data processing. The percentages of variance explained by the models are similar (around 70% of explained variance for drug sensitivity prediction) in the following three cases: unprocessed original data (thousands of genes), unprocessed original data for model-related genes only (tens of genes), and processed profiles of cell lines (tens of genes). This supports the choice of a model with a small number of relevant genes, which appear to contain most of the information needed for prediction. Second, the absolute level of performance appears much lower for CRISPR (between 30 and 50%) probably suffering from the lower number of samples, especially in cases where the number of variables is the highest. This tends to reinforce the interest of mechanistic approaches that do not use any training on the data for smaller datasets, less suitable for learning. Finally, while mutations and RNA data seem to provide the same predictive power (especially for drugs), using the two together does not necessarily result in a better performance.

Variable importance in these different random forests are reported in Fig S3 and are consistent with the analysis of mechanistic models. The mutational status of BRAF is definitely the most important variable followed by mutations in RAS or TP53. Concerning RNA levels, the most explanatory variables seem to be FOXD3 or PTEN, in line with model definitions.

## Discussion

The emergence of high-throughput data has made it easier to generate models for prognostic or diagnostic predictions in the field of cancer. The numerous lists of genes, biomarkers and predictors proposed have, however, often been difficult to interpret because of their sometimes uncertain clinical impact and little overlap between competing approaches [60]. Methods that can be interpreted by design, which integrate *a priori* biological knowledge, therefore appear to be an interesting complement able to reconcile the omics data generated and the knowledge accumulated in the literature.

These benefits come at the cost of having accurate expert description of the problem to provide a relevant basis to the mechanistic models. This is particularly true in this work since the personalized models all derive from the same structure of which they are partially constrained versions. It is therefore necessary to have a generic model that is both sufficiently accurate and broad enough so that the data integration allows the expression of the singularities of each cell line. If this is not the case, the learning of logical rules or the use of ensemble modelling could be favoured, usually including perturbation time-series data [61]. It should also be noted that, in the BRAF model presented here, the translation of biological knowledge into a logical rule is not necessarily deterministic and unambiguous. The choices here have been made based on the interpretation of the literature only. And the presence of certain outliers, i.e., cell lines whose behaviour is not explained by the models, may indeed result from the limitations of the model, either in its scope (important genes not integrated), or in its definition (incorrect logical rules). More global or data-driven approaches to define the model would be possible but would require different training/validation steps and different sets of data.

The second key point is the omics data used. For practical reasons, we have focused on mutation and RNA data. The legitimacy of the former is not in doubt, but their interpretation is, on the other hand, a crucial point whose relevance must be systematically verified. The omission or over-interpretation of certain mutations can severely affect the behaviour of personalized models. Validation using sensitivity data provides a good indicator in this respect. However, the question is broader for RNA data: are they relevant data to be used to personalize models, i.e., can they be considered good proxies for node activity? The protein nature of many nodes in the model would encourage the use of protein level data instead, or even phosphorylation levels if they were available for these data. One perspective could even be to push personalization to the point of defining different types of data or even different personalization strategies for each node according to the knowledge of the mechanisms at work in the corresponding biological entity. A balance should then be found to allow a certain degree of automation in the code and to avoid overfitting.

Despite these limitations, the results described above support the importance of combining the integration of different types of data to better explain differences in drug sensitivities. There was no doubt about this position of principle in general [62], and in particular in machine learning methods [3, 63]. The technical implementation of these multi-omic integrations is nevertheless more difficult in mechanistic models where the relationships between the different types of data need to be more explicitly formulated [8]. The present work therefore reinforces the possibility and value of integrating different types of data in a mechanistic framework to improve relevance and interpretation and illustrates this by highlighting the value of RNA data in addition to mutation data in predicting the response of cell lines to BRAF inhibition. In addition, one piece of data that could be further exploited is that of the specific behaviour of the drugs or inhibitors studied, since for instance some BRAF inhibitors have affinities that vary according to mutations in the BRAF gene itself. The integration of truly precise data on the nature of the drug is nevertheless limited by logical formalism and is more often found in less constrained approaches [64].

To conclude, we provide a comprehensive pipeline from clinical question to a validated mechanistic model which uses different types of omics data and adapts to dozens of different cell lines. This work, which is based only on the interpretation of data and not on the training of the model, continues some previous work that has already demonstrated the value of mechanistic approaches to answer questions about response to treatment, especially using dynamic data [65], and sometimes about the same pathways [8]. In this context, our approach proves the interest of logical formalism to make use of scarce and static data facilitating application to a wide range of issues and datasets in a way that is sometimes complementary to learning-based approaches.

## Supporting information

### Code

Code and data required to reproduce the study and all the plots are available in the following repositories:

- Model-checking: https://github.com/sysbio-curie/MaBoSS_test
- Data, model personalization and analyses: https://github.com/sysbio-curie/PROFILE_BRAR.Model

**Model** The model can be found in the dedicated GitHub: https://github.com/sysbio-curie/PROFILE_BRAF_Model/Model and will be deposited to model repositories in the logical section of BioModels database (http://biomodels.caltech.edu/) and in GINsim webpage (http://ginsim.org/models_repository) upon publication.

**S1 Fig. Mapping of the data in the RNAseq data on the regulatory network.** The expression data from both melanoma and colorectal cell lines used in this study are mapped onto the network. The scores correspond to the difference in the mean expression of the normalized data (using PROFILE method [32]). Red nodes show higher gene expression in melanomas and blue nodes higher expression in colorectal cancer cell lines. If most active nodes are equivalent to phosphorylated data, the mapping of RNAseq data informs on the gene status and the possibilities to activate the nodes. Thus, conclusions should be made with this fact in mind. At the gene level, then, genes such as SOX10, FOXD3, AKT, p21 and SPRY tend to have a higher expression in melanomas confirming their role in response to the treatment, whereas genes such as EGFR, ERBB2, MET, PTEN and FGFR2 are more relavant to colorectal cancers. This figure reinforces the idea that the mechanisms related to the response to anti-BRAF treatment may have different outcomes in both cancers bascule of a different gene context.

**S2 Fig. Performances of random forests for BRAF sensitivity prediction.** Random forests algorithms are trained with different omics types (mutations, RNA or both) and data processing (original data or processed data) to predict sensitivity to BRAF inhibition, through drug or CRISPR screening. Performances are expressed as percentage of explained variance by the fitted random forests

**S3 Fig. Variable importance for BRAF sensitivity prediction by random forests.** Variable importance for inhibition of BRAF by drugs (first row in Fig S2), when random forests algorithms are trained with different omics types (mutations, RNA or both) and data processing (original data or processed data). Higher values of variable importance correspond to higher decrease in prediction performance when the variable is disturbed by permutation and therefore to variables with a positive contribution to predictive performance.

## Acknowledgments

We would like to thank Celine Hernandez for fruitful discussions and inspiration during the elaboration of the verification tool, MaBoSS_test. LC, EB and VN acknowledge support from ANR in the context of the project ANR-FNR project AlgoReCell [ANR-16-CE12-0034]. This work was partially supported by the European Union’s Horizon 2020 program (grant 826121, iPC project). EB holds a chair at Paris Artificial Intelligence Research Institute Institute (PRAIRIE), funded in part by the French government under management of Agence Nationale de la Recherche as part of the “Investissements d’avenir” program, reference ANR-19-P3IA-0001 (PRAIRIE 3IA Institute).

