## Supplementary figures and images for "Personalized logical models to investigate cancer response to BRAF treatments in melanomas and colorectal cancers"

### Figure S1

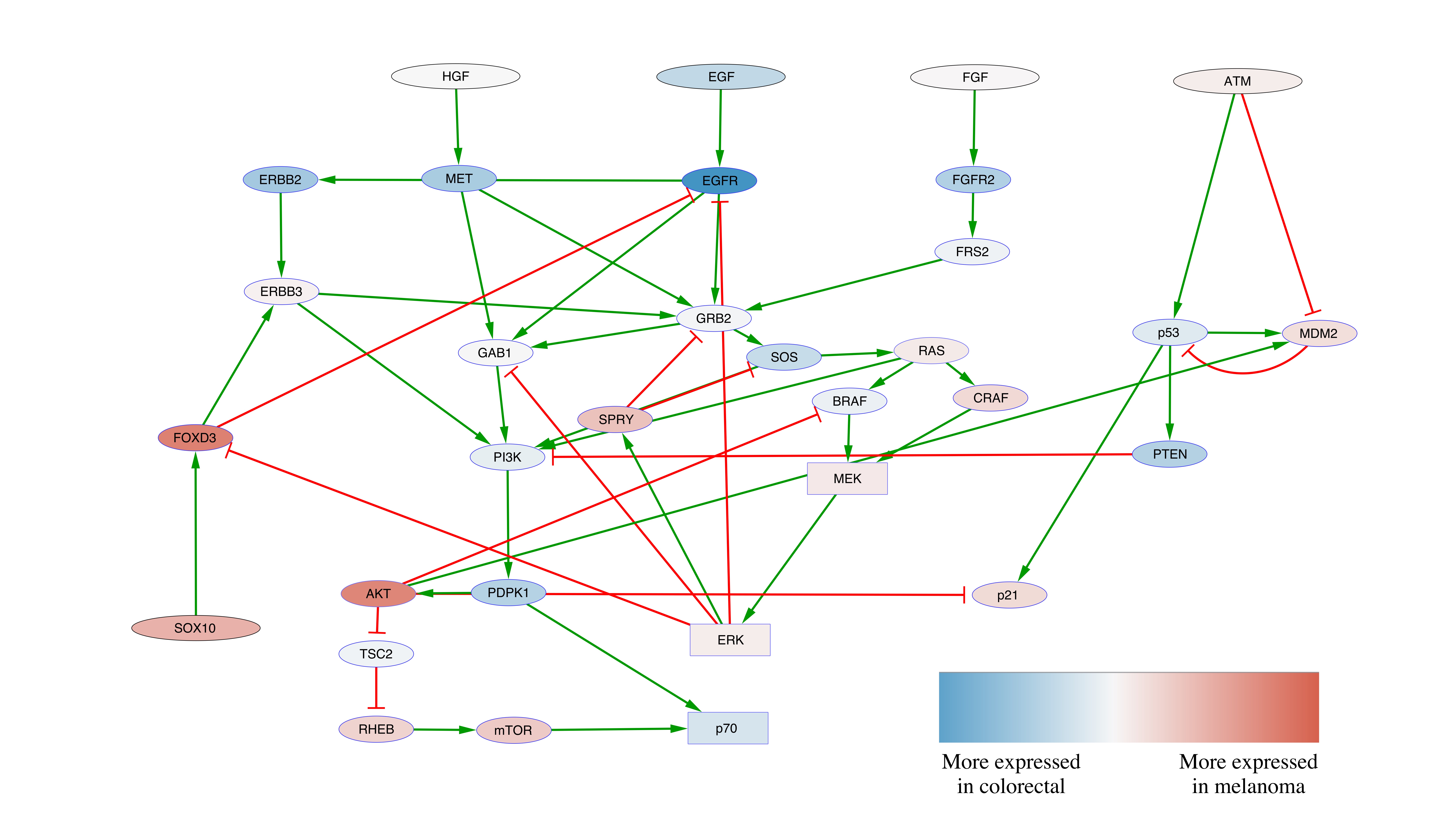

### Figure S2

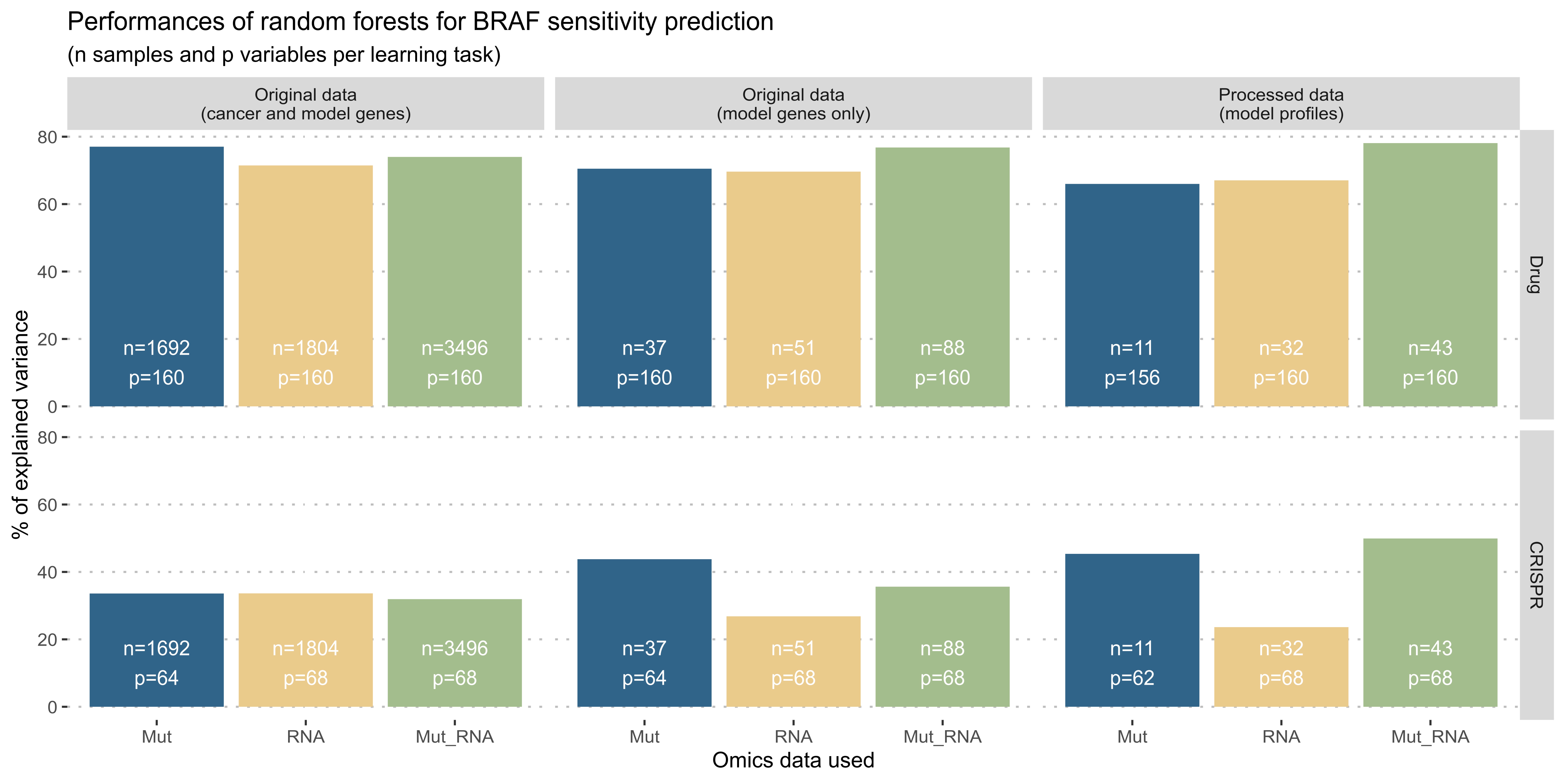

### Figure S3

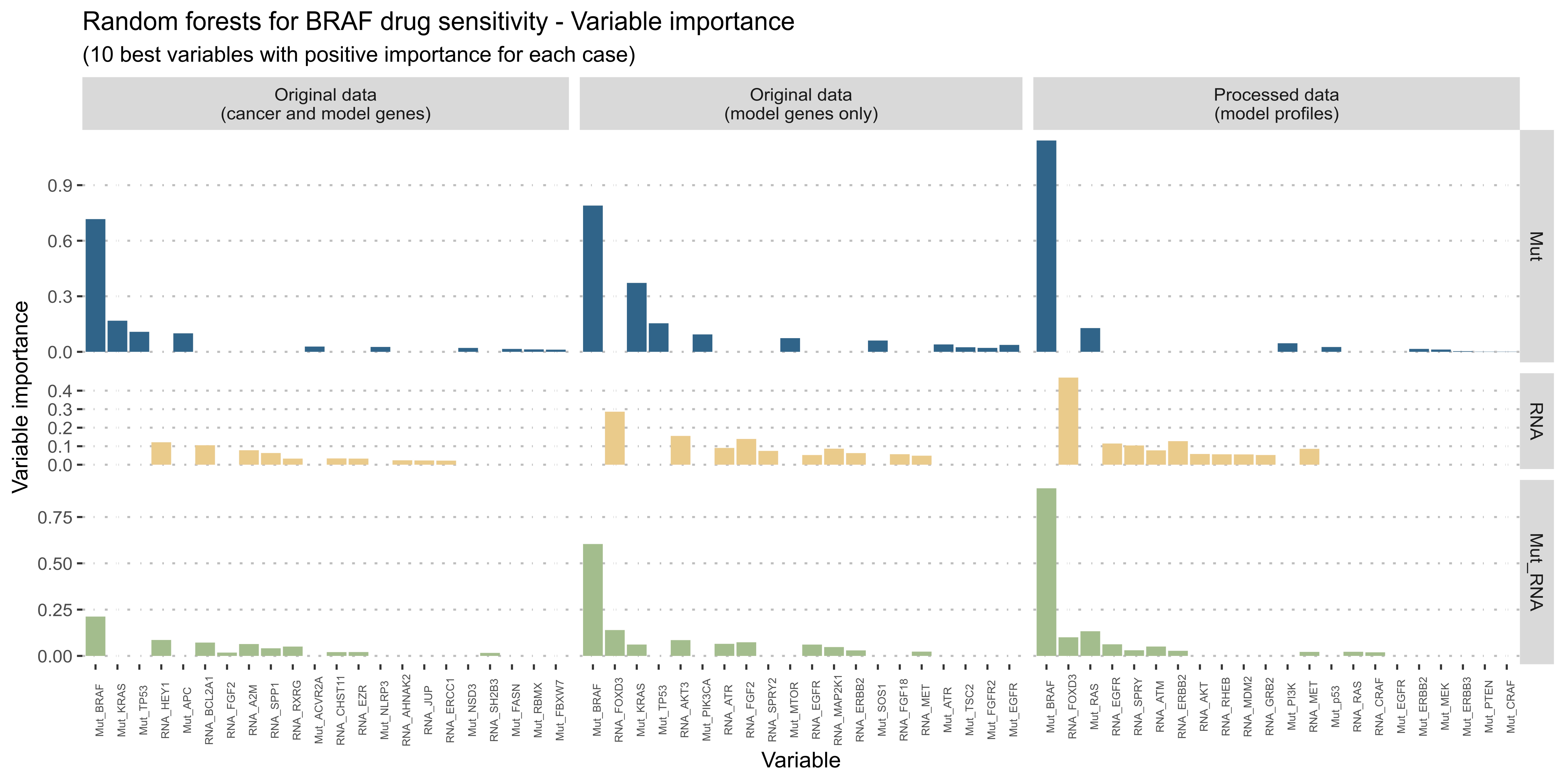
